# Mass mortality of foundation species on rocky shores: another reason why monitoring programs are relevant

**DOI:** 10.1101/2020.10.22.350769

**Authors:** María M. Mendez, Juan P. Livore, Federico Márquez, Gregorio bigatti

## Abstract

Global concern around substantial losses of biodiversity has led to the development of a number of large-scale long-term monitoring programs. In the past few decades, networks were established to obtain appropriate data on the spatial and temporal variation of marine species on rocky shores. Recently, the Marine Biodiversity Observation Network Pole to Pole of the Americas program (MBON P2P) was established and is coordinating biodiversity surveys along coastal areas throughout the continent. In this context, the goal of this paper was to demonstrate whether the proposed MBON P2P sampling protocol is capable of detecting rapid declines in cover of foundation species on Patagonian rocky shores. Changes in mussel beds cover were studied on monitored sites in northern Patagonia. Concurrently, long-term mussel bed dynamics were assessed based on existing data. Results showed that a mussel mortality event could be detected with this methodology. It took less than a year for mussel cover to drop from 90 to almost 0% despite the fact that significant changes in mussel bed cover were not registered in the previous 20 years at the study area. Therefore, yearly monitoring is needed, as a minimum, in order to timely perceive this kind of process. Real-time detection offers the opportunity of properly understanding the causes that lead to the loss of key community components such as these foundation species. Furthermore, it would provide early warning to decision makers enhancing the chances of conservation of natural environments and their key ecosystem services.

## Introduction

Rocky shores are one of the most widely distributed coastal habitat throughout the world (Thompson et al., 2002). Different human stressors, including introduction of species, physical modification of the coast, contamination, recreation, and the changes in climate are continuously threatening these habitats (Halpern et al. 2015; Duffy et al. 2019). Several efforts have been made to build coordinated international monitoring networks across the world aimed at obtaining temporal data on biodiversity, community structure and dynamics of rocky shores (Duffy et al. 2019; Canonico et al. 2019). Starting with the Marine Biodiversity and Climate Change Project (MarClim) in 1950 on the coast of United Kingdom and France, different programs have been developed with this premise around the world.

The history of monitoring networks in South America is relatively short; with three main efforts recorded in the last 20 years. The Natural Geography in Shore Areas (NaGISA) project of the Census of Marine Life (CoML) program (2000-2010) and its sequel the South American Research Group on Coastal Ecosystems (SARCE) (2009-2014), together monitored rocky shores over more than 60 degrees of latitude across 150 sites. Such monitoring described and analyzed biodiversity patterns across latitudinal gradients and linked them with ecosystem functioning and human stressors (Miloslavich et al., 2016; Cruz-Motta et al., 2020). Lastly in 2016, the Pole to Pole project of the Marine Biodiversity Observation Network (MBON P2P) was established with a goal of using common methods for the collection of biological information in coastal habitats throughout the American continent (Canonico et al., 2019; Duffy et al., 2019).

Argentinean Patagonia (41-55° S; 63-70° W) rocky intertidal shores were included as sampling sites since 2007 in the NaGISA and SARCE projects (Miloslavich et al., 2016; Cruz-Motta et al., 2020) and are currently being included in MBON P2P. One of the most important features on these shores is the extreme desiccation that intertidal organisms are exposed to through a combination of strong dry winds, low humidity and scarce rainfall (Bertness et al., 2006). Scorched mussel beds of the mid-intertidal are a distinctive component of the shores and a dense matrix of the two species, *Brachidontes rodriguezii* and *Perumytilus purpuratus,* dominate the physiognomy of the rocky shore communities (Bertness et al., 2006; Silliman et al., 2011, Miloslavich et al 2016). These communities are unique in that almost all intertidal organisms are unable to survive outside of the mussel bed, hence community structure and its diversity along with ecosystem function in these shores are obligately dependent on foundation species (Bertness et al. 2006). Historically, mussel beds from this region show simple structure, uniform appearance and disturbance-generated bare space throughout the bed is strikingly rare (Bertness et al., 2006; Adami et al., 2018). After the 2019 austral summer, losses in scorched mussels cover were visually observed at different monitored sites. This natural phenomenon gives us the opportunity of proving the usefulness and adequacy of a simple, low-cost, low-tech method proposed for the recently established MBON P2P program (Livore et al. under review). The goal of this paper was to demonstrate whether this sampling protocol is capable of detecting rapid declines in cover of foundation species on Patagonian rocky shores. Furthermore, we aimed to assess the stability and resilience of local mussel beds in order to describe their long-term natural dynamics.

## Methods

### Study sites and Sampling design

The study was performed at three rocky shores (Punta Cuevas, PC: 42°47’S; 65°00’W, Punta Este, PE: 42°47’S; 64°57’W and Punta Loma, PL: 42°48’S, 64°53’W) on the southwest coast of Golfo Nuevo, Chubut, Argentina. Tides at the sites are semidiurnal with mean amplitude of ~4 m, which exposes a sedimentary rock platform (consolidated mudstone). The characteristic three-level biological zonation of Patagonian rocky shores is present at the sites studied. The high intertidal zone has a large proportion of bare rock, being the invasive barnacle *Balanus glandula* and the limpet *Siphonaria lessonii* abundant. The mid-intertidal is dominated by matrix of scorched mussels, *Brachidontes rodriguezii* and *Perumytilus purpuratus.* The low intertidal level is characterized by several ephemeral algal species and a large proportion of the calcareous alga *Corallina officinalis* (Bertness et al., 2006; Miloslavich et al., 2016). PC and PE were chosen as a monitoring sites of the SARCE project and the three sites are currently part of the MBON program.

Scorched mussel cover data before (2014 and 2018) and after (2019) the observation of scorched mussel beds declines (austral summer 2019) were compared for the three sites. A specifically designed protocol to study changes in rocky shore communities was used in all monitoring events (adapted from Ocean Best Practices: http://dx.doi.org/10.25607/OBP-5). Briefly, percentage cover of mussel bed was estimated inside 25 x 25 cm photoquadrats haphazardly placed on the substrate (n = 5 to 10 per sampling; N = 50). Recently, this method was compared to a previously used *in situ* visual method and both similarly detected spatial and temporal variability of rocky shores assemblages (Livore et al. under review). In the lab, one hundred equidistant points were placed over the digital image using the free software Coral Point Count (CPCe V 4.1, Kohler and Gill, 2006) and organisms observed under each point were determined to the lowest possible taxonomic level. Statistical differences in mussel cover between sites for the before samplings were evaluated using a one-way ANOVA. Normality and homogeneity of variance assumptions were evaluated with Kolmogorov-Smirnov and Levene tests, respectively. Percentage cover of mussels in the 2019 samplings was close to zero, thus this year was not statistically compared with the previous samplings.

To describe the long-term natural dynamics of local mussel beds we collected historical data (e.g. cover) from Punta Cuevas. This intertidal is located at the southern end of the city of Puerto Madryn, 500 meters from the CCT CONICET-CENPAT and the Universidad Nacional de la Patagonia (the authors’ workplaces). Both scientist and biology professors, from these institutions, regularly use the site for academic and scholar field trips. Thus, quantitative and qualitative (for example, panoramic photographs) data of mussel cover and abundance was obtained almost annually for the last 20 years to examine the stability and resilience of the beds (Table 1).

**Table 1.**
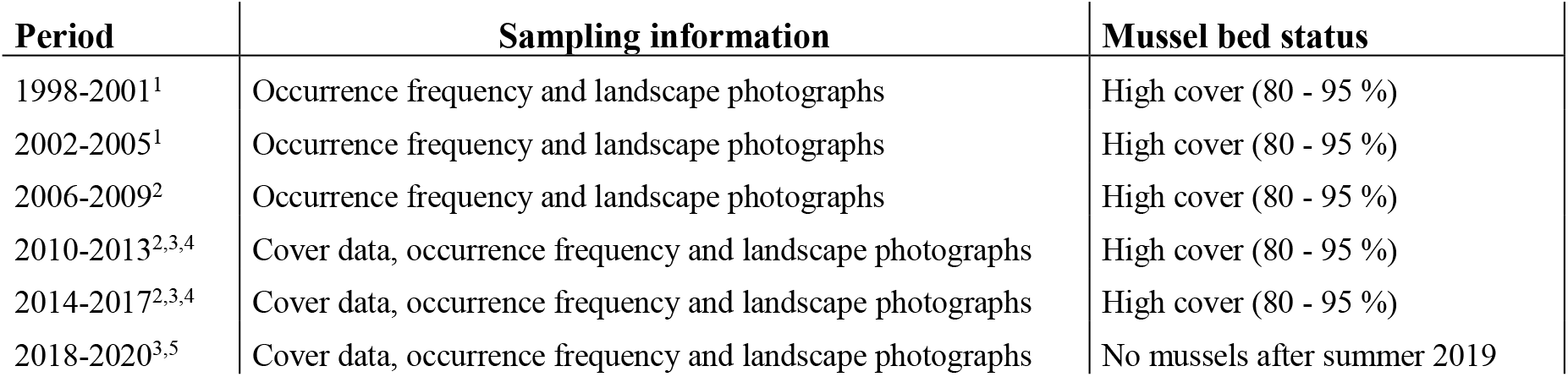
Natural dynamics of Punta Cuevas mussel bed between 1998 and 2020. Sources included: ^1^ Torres and Caille 2009, ^2^ “Community ecology” (Prof. A. Bisigato, UNPSJB) samplings, ^3^ “Malacology” (Prof. G. Bigatti, UNPSJB) samplings, ^4^ SARCE samplings, ^5^ MBON samplings and a vast photographic record of the site

| Period | Sampling information | Mussel bed status |
| --- | --- | --- |
| 1998-2001<sup>1</sup> | Occurrence frequency and landscape photographs | High cover (80 - 95 %) |
| 2002-2005<sup>1</sup> | Occurrence frequency and landscape photographs | High cover (80 - 95 %) |
| 2006-2009<sup>2</sup> | Occurrence frequency and landscape photographs | High cover (80 - 95 %) |
| 2010-2013<sup>2,3,4</sup> | Cover data, occurrence frequency and landscape photographs | High cover (80 - 95 %) |
| 2014-2017<sup>2,3,4</sup> | Cover data, occurrence frequency and landscape photographs | High cover (80 - 95 %) |
| 2018-2020<sup>3,5</sup> | Cover data, occurrence frequency and landscape photographs | No mussels after summer 2019 |

## Results and Discussion

This study shows how a simple low-cost and non-extractive method was able to provide real-time data on changes in rocky shores biodiversity, as a mussel mortality event and served as a sentinel of a process occurring along Atlantic Patagonian rocky shores. Anthropogenic pressure on coastal areas is increasing worldwide and detecting temporal and spatial changes in rocky shore biodiversity is critical for its conservation (Duffy et al., 2019). Those sites suffering significant changes in biodiversity are considered degraded or unhealthy ecosystems. This equates to risks to different ecosystem services for billions of people, some as essential as human nutrition and health, recreation attractions or public safety (Canonico et al., 2019). One way to detect biodiversity changes is the implementation of monitoring programs. Although the history of these programs is quite young in South America, the present study demonstrates the efficacy of their methodologies in detecting rapid biodiversity changes. In the study area, dense beds of scorched mussels are a distinct component and the foundation species in the mid-intertidal (Bertness et al., 2006; Adami et al., 2018; Miloslavich et al 2016). Mussel bed cover at different sites of Golfo Nuevo was close to 90% before the 2019 austral summer (no differences were detected between sites: F = 0.46, df = 2, p = 0.63. Fig. 1). However, almost no scorched mussels were registered at the sites after the 2019 summer (Fig. 1).

**Fig. 1.**
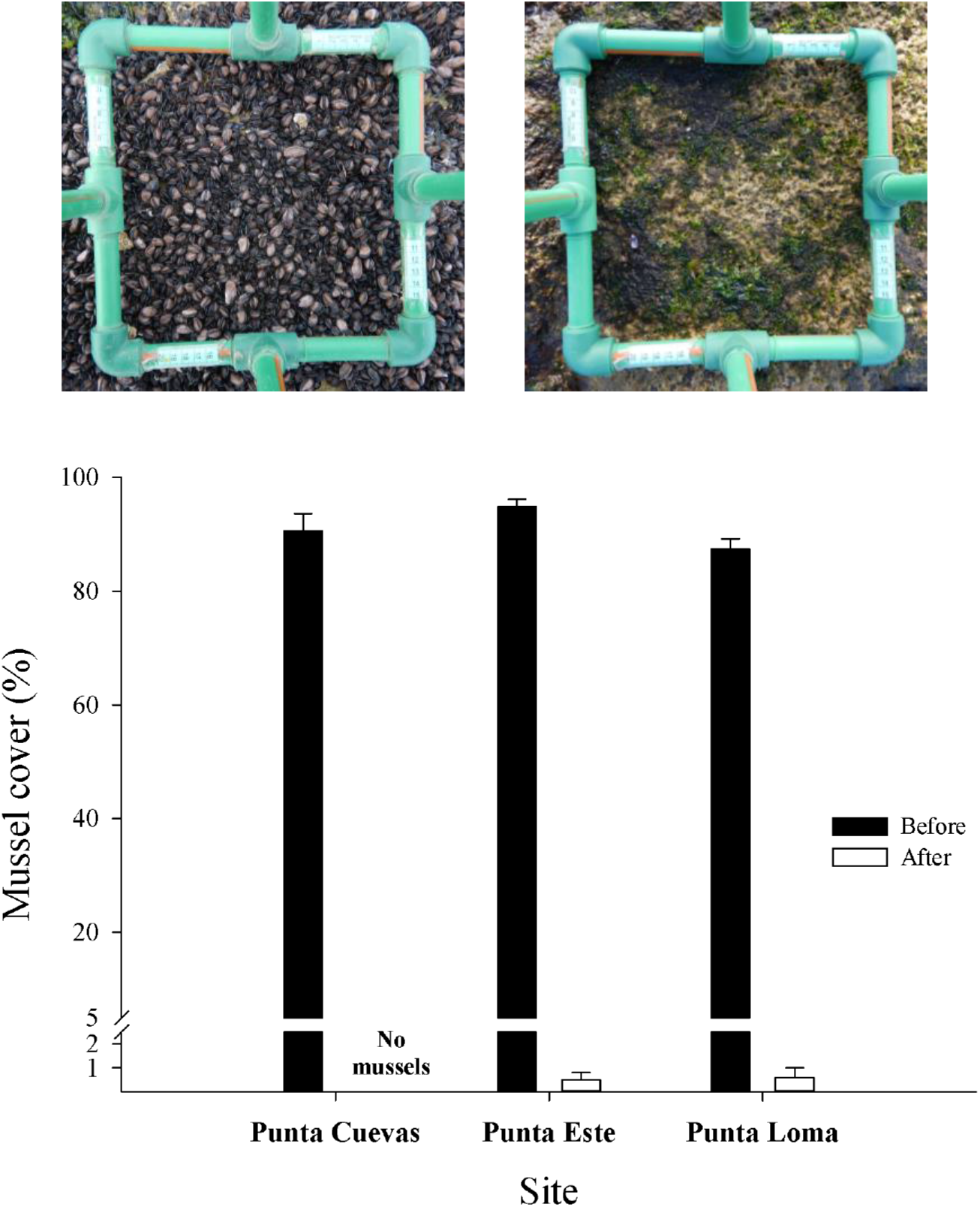
Top: Plots in the mid intertidal of Punta Cuevas showing the natural cover of the mussel bed in October 2018 (left) and after the mass mortality event, when mussels were absent on all horizontal surfaces in June 2019 (right). Bottom: Mean (SE) mussel cover for the three samplings sites before and after the mortality event

In northern Patagonian rocky shores desiccation plays a fundamental role in community structure (Bertness et al., 2006). Organisms are exposed to strong dry winds, combined with low humidity and scarce rainfall (Bertness et al., 2006). Only a few habitat-forming species are capable of tolerating these extreme conditions. In this sense, mussels act as ecosystem engineer species and provide habitats and refuge for other organisms (Silliman et al., 2011). More than 40 invertebrate species live associated with mussel beds avoiding environmental stress (Silliman et al., 2011). Thus, the observed loss of the mussel cover would have important indirect effects on the associated assemblage through the absence of tens of species.

Our examination of the natural dynamics of Punta Cuevas mid-intertidal showed that significant changes in cover and abundance of scorched mussel are extremely uncommon (Table 1). Beds at the site are highly stable and resistant to disturbance, as was suggested by Bertness et al. (2006) for central Patagonian rocky shores. During the last 20 years, no visible change of mussel cover was registered at the study site in spite of being under high anthropogenic (e.g. trampling) and natural disturbance pressure (Mendez et al. 2019). Furthermore, photographic records show that bare rock patches never exceeded ~10 m^2^ in the mussel bed during this period. This contrasts with mussel beds on the Atlantic and Pacific coasts of North America, for example, where disturbance-generated patches are common (Paine and Levin, 1981). The registered stability could be related to the small size of scorched mussels (usually about 0.20 cm long) and its strong byssal attachment strength (Bertness et al. 2006; Mendez et al. 2019). Altogether, our results suggest that that the drastic decline of scorched mussel cover observed at the study sites were not a consequence of the natural dynamics of the habitats and that this event was not part of the baseline fluctuations of the mid rocky shore community.

From a conservation perspective, it is essential to consider the recovery time needed by a given community to reestablish after a disturbance like the one reported here. Although *B. rodriguezii* has been observed to recruit continuously during the year (Arribas et al., 2015) in northern Atlantic Patagonia, it would take up to a decade to reach the full recovery of the mussel bed (Bertness et al. 2006; Mendez et al. 2017). After the mass mortality event, bare rock became available for colonization by early successional organisms, such opportunistic algae and barnacles. The switch in species dominance at the sites (i.e. barnacles and algae replacing mussels) could last several years and would influences local biological interactions with important consequences for community structure. Thus, in systems exposed to high physical stress where foundation species are dominant, management and conservation efforts should be focused in the foundation species instead on more charismatic organisms that live associated to them (Bertness et al. 2006).

When massive mortalities occur, there is a need to study the causes of the event. Fungi, parasites, bacteria, viruses or toxic blooms have been reported as responsible agents for this kind of events in bivalves (Peperzak and Poelman, 2008; Vásquez et al. 2016). More recently, several studies documented how extreme weather events (e.g. heatwaves) can be responsible for mass mortalities in dominant organisms (Zamir et al., 2018), including mytilids (Seuront et al., 2019). At the time of publication, potential causes for this sudden mass mortality are being studied and none of them could be completely ruled out.

This work showed how quick the mass mortality event happened, taking less than 6 months for the mussel beds to decline from 90% to losing all of its cover (Fig. 1). Significant effort is spent on debating sampling frequency in monitoring programs; our results contribute empiric evidence that highlights the need to have at least yearly monitoring. It is important to note that without periodic monitoring, this mortality case would not have been detected in its beginnings and the causes that led to it could not have been studied properly. The immediacy of the detection gives us the opportunity to study the changes in the community from the start and to follow the entire recovery process, whilst considering the concomitant effects on ecosystem function. Studies of this nature, derived from monitoring programs, can give early alarm to decision makers and provide a timely response action that mitigates the impacts on coastal zones and preserve rocky shore environments.

## Acknowledgements

We thank Pepe Ojeda for technical support. Agencia Nacional de Promoción Científica y Tecnológica (PICT 2018-0969) partially support this study. This is publication #XX of the Laboratorio de Reproducción y Biología Integrativa de Invertebrados Marinos (LARBIM).

## References

Adami, M., Schwindt, E., Tablado, A., Calcagno, J., Labraga, J. C., & Orensanz J.M. (2018). Intertidal mussel beds from the South-western Atlantic show simple structure and uniform appearance: does environmental harshness explain the community? Marine Biology Research, 14, 403–419.

Arribas, L.P., Bagur, M., Gutiérrez, J. L., & Palomo, M. G. (2015). Matching spatial scales of variation in mussel recruitment and adult densities across southwestern Atlantic rocky shores. Journal of Sea Research, 95, 16–21.

Bertness, M. D., Crain, C. M., Silliman, B. R., Bazterrica, M. C., Reyna, M. V., Hidalgo, F. et al. (2006). The community structure of Western Atlantic Patagonian rocky shores. Ecological Monograph, 76, 439–460.

Canonico, G., Buttigieg, P. L., Montes, E., Muller-Karger, F. E., Stepien, C. A., Wright, D. et al. (2019). Global observational needs and resources for marine biodiversity. Frontiers in Marine Science, 6, 367.

Cruz-Motta, J. J., Miloslavich, P., Guerra-Castro, E., Hernández-Agreda, A., Herrera, C., Barros, et al. (2020). Latitudinal patterns of species diversity on South American rocky shores: local processes lead to contrasting trends in regional and local species diversity. Journal of Biogeography, https://doi:10.1111/jbi.13869

Duffy, J. E., Benedetti-Cecchi, L., Trinanes, J., Muller-Karger, F. E., Ambo-Rappe, R., Boström, C. et al. (2019). Toward a coordinated global observing system for seagrasses and marine macroalgae. Frontiers in Marine Science, 6, 317.

Halpern, B. S., Frazier, M., Potapenko, J., Casey, K. S., Koenig, K., Longo, K. et al. (2015). Spatial and temporal changes in cumulative human impacts on the world’s ocean. Nature Communications, 6, 7615.

Kohler, K. E., & Gill, S. M. (2006). Coral Point Count with Excel extensions (CPCe): a Visual Basic program for the determination of coral and substrate coverage using random point count methodology. Computers and Geosciences, 32, 1259–1269.

Livore, J. P., Mendez, M. M., Miloslavich, P., Rilov, G., & Bigatti, G. Biodiversity monitoring in rocky shores: challenges of devising a globally applicable and costeffective protocol. Ocean and Coastal Management (under review).

Mendez, M. M., Livore, J. P., Calcagno, J. A., & Bigatti, G. (2017). Effects of recreational activities on Patagonian rocky shores. Marine Environmental Research, 130, 213–220.

Mendez, M. M., Livore J. P., & Bigatti G. (2019). Interaction of natural and anthropogenic stressors on rocky shores: community resistance to trampling. Marine Ecology Progress Series, 631, 117–126.

Miloslavich, P., Cruz-Motta, J. J., Hernandez, A., Herrera, C. A., Klein, E., Barros, F. et al. (2016). Benthic assemblages in South American intertidal rocky shores: biodiversity, services, and threats. In: R. R. Rodríguez (Ed.) Marine Benthos: Biology, Ecosystem Functions and Environmental Impact. New York: Nova Science Publishers.

Paine, R. T., & Levin S. A. (1981). Intertidal landscapes: disturbance and the dynamics of pattern. Ecological Monographs, 51, 145–178.

Peperzak, L., & Poelman, M. (2008). Mass mussel mortality in The Netherlands after a bloom of *Phaeocystis globosa* (Prymnesiophyceae). Journal of Sea Research, 60, 220–222.

Seuront, L., Nicastro, K. R., Zardi, G. I., & Goberville, E. (2019). Decreased thermal tolerance under recurrent heat stress conditions explains summer mass mortality of the blue mussel *Mytilus edulis*. Scientific Reports, 9, 1–14.

Silliman, B. R., Bertness, M. D., Altieri, A. H., Griffin, J. N., Bazterrica, M. C., Hidalgo, F. J. et al. (2011). Whole-community facilitation regulates biodiversity on Patagonian rocky shores. PLoS ONE 6, e24502.

Thompson, R. C., Crowe, T. P., & Hawkins, S. J. (2002). Rocky intertidal communities: past environmental changes, present status and predictions for the next 25 years. Environmental conservation, 168

Torres, A., & Caille, G. (2009). Las comunidades del intermareal rocoso antes y después de la eliminación de un disturbio antropogénico: un caso de estudio en las costas de Puerto Madryn (Patagonia, Argentina). Revista de Biología Marina y Oceanografía, 44, 517–521.

Vázquez, N., Fiori, S., Arzul, I., Carcedo, C., & Cremonte, F. (2016). Mass mortalities affecting populations of the yellow clam *Amarilladesma mactroides* along its geographic range. Journal of Shellfish Research, 35, 739–745.

Zamir, R., Alpert, P, & Rilov, G. (2018). Increase in weather patterns generating extreme desiccation events: implications for Mediterranean rocky shore ecosystems. Estuaries and Coasts, 41, 1868–1884.

